# Microscope control with a natural language agent

**DOI:** 10.64898/2026.09.15.751723

**Authors:** Zach Marin, Amr Abouelezz, Nestor Miguel Castillo Duque de Estrada, Sarah V. Schweighofer, Fabian Hauser, Leia T. Köstinger, Anvita Manjunath, Nico Stuurman, Florian Schueder, Jonas Ries

## Abstract

Microscopy underpins modern biology, but acquiring high-quality data from complex experiments at scale remains challenging. Smart microscopy can standardize and automate data collection, but each experiment-specific solution requires coding expertise and development time. To address this, we present MicroClaw, an AI agent that advises on experiment- and system-specific parameters and approaches and collaboratively plans and executes diverse, complex, and reusable imaging workflows across microscope platforms, without requiring expert knowledge.

## Introduction

Microscopy is a fundamental experimental pillar of modern biological research, shedding light on the spatial organization of biological systems from entire tissues down to individual biomolecules. Given its enormous range of applications across cell types, organisms, and virtually all disciplines of biology, it is not surprising that the requirements for image acquisition vary dramatically between experiments. In many cases, optimal imaging conditions can become highly complex and depend on the biological sample, the process being observed, and the specific information one aims to extract. It can be challenging and laborious to execute repeated imaging experiments that meet these conditions. To address this, researchers have built smart microscopes^1,2^ to automatically identify relevant biological features or events and acquire data on a user’s behalf, accelerating existing workflows and enabling new ones that are too long for users to execute manually. However, relatively few experimental microscopy tasks have been automated with smart microscopy due to the software engineering challenges of dynamically controlling hardware in response to acquired data^2^.

To alleviate the software engineering burden of smart microscopy, recent studies and patents have proposed using large language model (LLM) agents to convert natural language user input into actions taken on laboratory equipment^3^, including on fluorescence and atomic force microscopes^4–8^. In addition to their programming capabilities, LLMs should be capable of acting as collaborative scientists when imaging, offering suggestions on parameter optimization and troubleshooting when the instrument malfunctions. While putting LLMs on microscopes is a logical next step in the rapidly developing field of AI, the plethora of laboratory equipment and experimental tasks^2^ makes it challenging to develop a general AI-guided hardware assistant for research.

Here, we develop an LLM agent, which we call MicroClaw, that uses Claude (Anthropic, San Francisco, CA, USA) and Micro-Manager^9,10^ to control most hardware currently used in microscopy-based research. This powerful combination enables MicroClaw to (1) guide users in selection of parameters and settings, automatically optimize these per sample and per question, and troubleshoot common acquisition problems and (2) collaboratively develop smart microscopy routines that deterministically automate time-consuming tasks, such as imaging large sample areas or identifying rare events. Together, these capabilities enable users to take better data and implement automated, complex workflows that are challenging to perform manually.

## Results

### Microclaw architecture

We developed MicroClaw as a pseudo-agentic^11^ tool capable of coordinating microscopy hardware, developing and using analysis routines, integrating external analysis software, and combining these in acquisition feedback loops. It is easy to set up, featuring a single-click installer and a first launch interview to discuss how researchers expect the microscope to work, including clarification of any uncertainties from the perspective of the agent (e.g. where in the optical path are each of the two filter wheels). It operates in parallel with Micro-Manager, allowing users to either control the microscope directly through the Micro-Manager interface or via conversation with the agent in a web-based GUI (**Figure 1a**). MicroClaw connects to Micro-Manager via Pycro-Manager^12^, a Python interface for Micro-Manager. It can connect and disconnect from Micro-Manager seamlessly, operating only when it is needed. Furthermore, MicroClaw learns about the microscope setup, experimental workflows, and individual user preferences, and stores this information in a persistent knowledge base on each system, which is retrieved at the start of each MicroClaw session.

**Figure 1:**
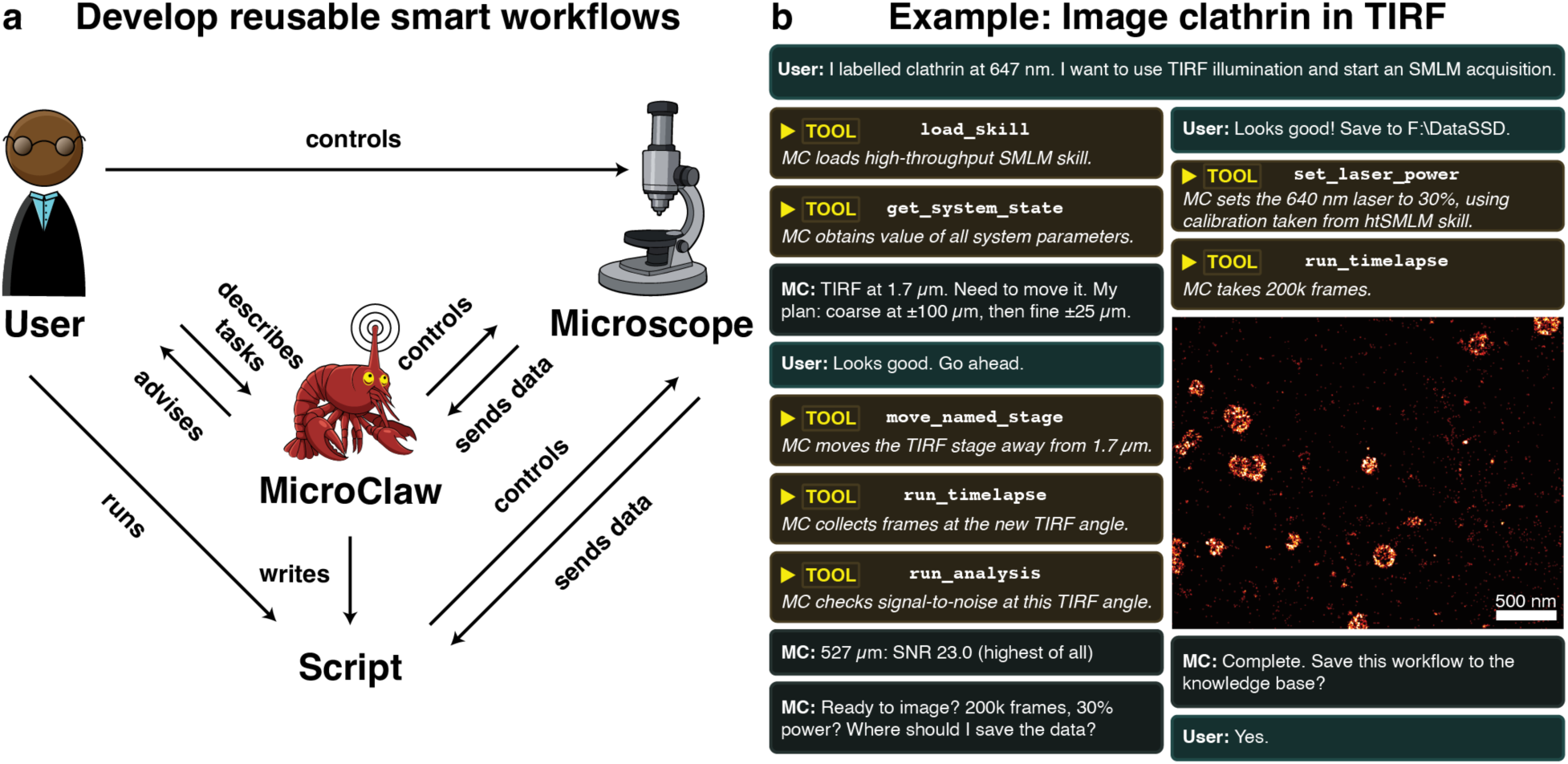
**a, MicroClaw-User workflow.** The user is able to interact with MicroClaw and/or the microscope independently. If the user passes a request to the MicroClaw agent, it is converted into a series of tool calls on the microscope. If the agent is unclear on the task, it asks the user for clarification. As the agent executes the request on the microscope, changes are reflected in the Micro-Manager GUI. Once the agent has executed the resulting set of tools, it passes the results back to the user. If the user is satisfied with the process, they can request the agent to export the sequence of tool calls to a Python script that runs independently of the agent. At this point, the agent can be closed and the user can execute the script repeatedly without any calls to the agent. **b, A visualization of a small example using MicroClaw**. The user asks MicroClaw (MC) to take an SMLM image of a clathrin sample. The agent converses with the user and calls tools as needed, resulting in actions on the microscope and an image (Figure 2a**, Supplementary Figure 1a**) with nanometer resolution.

Simple actions on the microscope are performed using “tools”, which are Python functions included with MicroClaw that move microscope stages, directly interact with the Micro-Manager GUI, save and load images, and so on. For acquisition tasks not covered by existing tools, MicroClaw writes custom Python functions, called “hooks,” which execute the user’s desired behavior.

To process image data, MicroClaw can write classical image analysis routines into hooks using scikit-learn^13^. These routines can be used to make simple image-based decisions in the acquisition pipeline. For more complicated image analysis, we provide connections to ilastik^14^, and it is possible to patch other existing and external analysis software into hooks.

Once a workflow, which may include both pre-built tools and user-defined hooks, has been developed, users can export it to a Python script and run this without requiring further interaction with the agent (**Supplementary Note 1**). Thus, a non-deterministic LLM session can yield deterministic acquisition scripts that can be used for experiments and published.

Approaches to developing analysis and acquisition workflows, and to troubleshooting, are stored in MicroClaw as “skill” files. MicroClaw skills include instructions for building hooks, for the proper sequencing of tool calls when acquiring single-molecule localization microscopy^15–18^ (SMLM) images, and for troubleshooting the optical path when no signal is detected by the camera. These are automatically loaded when relevant to the MicroClaw session.

### Simple microscope control

An example session demonstrating development of a simple MicroClaw workflow is shown in **Figure 1b**. Here, the user requests that MicroClaw take an image of a clathrin sample. The agent breaks this down into smaller tasks and provides its thinking and feedback, both as descriptive text and as tool calls that are displayed to the user as they are run. MicroClaw identifies the optimum TIRF angle and images **(Figure 2a, Supplementary Figure 1a**) using its built-in tools. This procedure can then be exported to the knowledge base and to a Python script, which is a reusable, deterministic repeat of a session (**Supplementary Note 1**).

**Figure 2:**
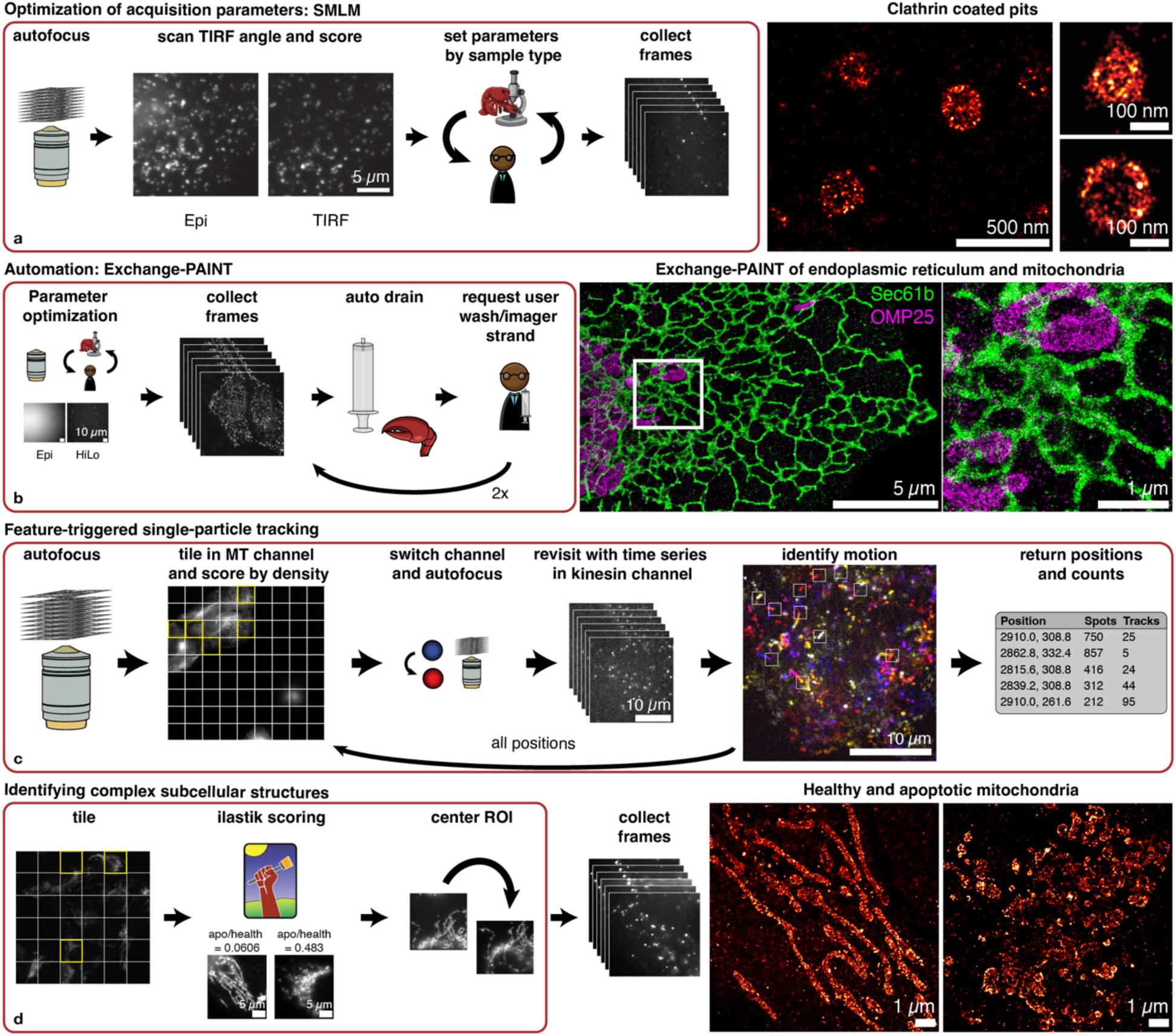
MicroClaw workflows, with agent-controlled areas boxed in red. **a, Optimization** of SMLM acquisition parameters on custom SMLM microscope for imaging of endocytic structures. MicroClaw optimized TIRF angle, set up an SMLM imaging sequence, and acquired images (clathrin light chains CLCa- and CLCb-SNAP labeled with BG-AF647). **b, Automated Exchange-PAINT** on a custom Nikon Eclipse Ti system. Here, the agent informed the user when to add new washes and imager strands. MicroClaw handled the rest. DNA-PAINT imaging of endoplasmic reticulum (green, Sec61b-mCherry, mCherry anti-mouse nanobody with a 5xR4 sequence and the R4 imager with a Cy3b dye at the 3’ end) and mitochondria (magenta, OMP25-GFP with an anti-GFP nanobody with a 5xR2 sequence, where the R2 imager has a Cy3B dye conjugated to the 3’ end). **c, MicroClaw screened for walking kinesins** on a coverslip, which were labeled in live cells expressing Kif5B[1-560]-HaloTag7 labeled with JFX646-HTL. MicroClaw identified and focused on the microtubules (expressing eGFP-ɑTubulin), then autonomously switched to a second channel to identify walking kinesins, their density and motility. **d, Automatic imaging of complex mitochondria structures** identified with machine learning software. MicroClaw was used to identify healthy and apoptotic mitochondria (both labeled with anti-TOMM20-AF488 for widefield and anti-TOMM20-AF647 for SMLM) on a coverslip using a pre-trained Ilastik^14^ model. Fields of view containing the desired structures were centered automatically. We used the Micro-Manager multi-dimensional acquisition wizard and the htSMLM plugin^25^ to collect SMLM data of the resulting FOVs.

### Optimization of acquisition parameters

MicroClaw can optimize acquisition parameters based on its knowledge of the microscope parameters, sample type (provided a relevant skill file is available), inspection of images, and interaction with the user.

As a test of parameter optimization capabilities, we asked MicroClaw to assist us in taking SMLM images of clathrin on one of our custom microscopes^19,20^ in total internal reflection fluorescence microscopy (TIRF) mode. It is desirable to operate in TIRF to focus on membrane-associated clathrin structures and minimize background from intracellular, out of focus labels. We asked MicroClaw to find the optimum TIRF angle and then start a STORM acquisition. MicroClaw figured out how to optimize the TIRF angle automatically, using four imaging metrics (**Supplementary Video 1, Supplementary Figure 1a**), and moved the stage into the correct position. Then, using information from an SMLM skill we added to MicroClaw, and feedback from the user, we identified ideal laser powers and exposure times and acquired a super-resolved image of clathrin (**Figure 2a**).

Parameter optimization was again helpful for imaging DNA origami^21^ on an Andor Dragonfly microscope. MicroClaw developed a second TIRF optimization procedure, similar to that for clathrin, for imaging samples with DNA points accumulation for imaging in nanoscale topography^22,23^ (DNA-PAINT), which is a subtype of SMLM. A DNA-PAINT skill, which was distilled from popular papers^15,24^ and our general knowledge, then helped the agent choose parameters for acquisition. The average-based reconstruction of the origamis (**Supplementary Figure 3**) is shown in **Supplementary Figure 1b.** MicroClaw was able to achieve sub-nanometer localization precision (0.9 nm) on an average of a 5-nm spaced custom-built DNA origami lobster (**Supplementary Figure 1b**).

### Automated microscopy

Another feature of MicroClaw is its ability to automate long and repeated acquisitions. Exchange-PAINT^26^ imaging can be a time-consuming procedure due to the sequential nature of the approach. To automate this process, we connected a syringe pump to a custom Nikon-Eclipse Ti system (see Methods), asked MicroClaw to perform sequential multiplexed imaging and informed it that the syringe pump could be used to automatically drain the system. MicroClaw inspected the available syringe-pump controls, asked for the syringe diameter to calculate accurate flow volumes, and subsequently constructed a workflow. It acquired images, automatically drained the imaging chamber with the pump when an Exchange-PAINT round was done, informed us to then add a wash and an imager strand, and then started the next round of imaging automatically. We used MicroClaw to acquire a two-round Exchange-PAINT image of mitochondria (OMP25-GFP with an anti-GFP nanobody with a 5xR2 sequence, where the R2 imager has a Cy3B dye conjugated to the 3’ end) and the endoplasmic reticulum (Sec61b-mCherry, mCherry anti-mouse nanobody with a 5xR4 sequence and the R4 imager with a Cy3b dye at the 3’ end) (**Figure 2b, Supplementary Figure 1b**).

This illustrates how existing hardware ecosystems can be rapidly incorporated into MicroClaw-assisted workflows. We expect that MicroClaw could automate the entire Exchange-PAINT workflow if we provided it with a more capable fluidics setup. In the context of the rapidly developing field of spatial biology^26–31^, such functionality could facilitate the implementation of automated fluidics pipelines^32^.

### Smart microscopy

MicroClaw is able to convert natural language descriptions of a task into reusable code that can make acquisition decisions based on collected images. We tested this capability by identifying subcellular regions in living cells containing walking kinesins. For these tracking experiments, we used a custom widefield microscope^19^ to find regions that contained both a clear microtubule network and well-separated walking trajectories of human kinesins (expressing Kif5B[1-560]-HaloTag7 labeled with JFX646-HTL). MicroClaw scanned and mapped large areas of the coverslip to recognize fields of view (FOVs) that contained fluorescent microtubules (expressing eGFP-ɑTubulin). It developed and applied a hook that identified flat microtubule networks near cell edges. It then switched laser wavelength and dichroic filter to look for kinesins in the detected cell periphery, and to assess their density based on the number of detected punctae. It found the closest plane where most kinesins were in focus and, if a good kinesin density was observed, MicroClaw marked those positions and proceeded to acquire time-lapse data. Next, MicroClaw identified walking kinesin given a threshold of brightness and speed of the punctae, and created new images of the timelapse with yellow boxes overlaid on the moving kinesins in each frame (**Figure 2c, Supplementary Figure 1c**). This will enable us in the future to automatically compare kinesin motilities across cell lines and conditions.

We next wanted to image specific regions that only a complex machine learning algorithm can identify. In order to study the effects of apoptotic proteins on mitochondria, cells with mitochondria in an adequate apoptotic state need to be found. We decided to use ilastik^14^ to identify healthy vs. apoptotic mitochondrial networks on a coverslip imaged on our custom SMLM system^20^ (**Figure 2d, Supplementary Figure 1d**). We first trained the ilastik network on example images of mitochondria labeled with anti-TOMM20-AF488. Then, we pointed MicroClaw to the location of the ilastik file. After automated acquisition of a tiled grid on the coverslip, MicroClaw scored each of the acquired FOVs in ilastik using the provided file. It correctly identified regions that contained mostly apoptotic or healthy mitochondria, depending on the user request. Next, MicroClaw was asked to center these mitochondria in the FOV and save a position list of these FOVs. We were then able to take SMLM images of the same mitochondria, which were also labeled with anti-TOMM20-AF647, in each FOV. This will allow us to identify cells in different apoptotic stages, and identify apoptotic mitochondria in mixed populations of cells or in tissues.

### Troubleshooting

MicroClaw is aware of all of the microscope settings available to Micro-Manager and comes with an optical layout skill. As such, when unforeseen problems arise, MicroClaw is able to identify and discuss possible solutions with the user. As an example, we used it to identify the source of image duplication on our camera, which was a magnetically mounted dichroic mirror in our ratiometric imaging path^19^ (**Supplementary Note 2**). Because MicroClaw cannot know if there is something physically in the light path that is not computer-controlled, the user was asked to have a look inside of the microscope to verify the existence of and remove the mirror. Even with its partial world view, MicroClaw correctly identified the source of the image duplication and guided the user in its removal.

## Discussion

Here, we demonstrated that MicroClaw can guide users in parameter selection, optimize parameters directly, troubleshoot imaging problems, and assist with or autonomously execute diverse microscopy acquisition workflows across multiple microscope platforms. Because users communicate with MicroClaw via natural language, no coding experience is required.

We use foundation models, in particular Claude Opus, to achieve this capability, and attempt to address challenges that come with their use: lack of data privacy, cost, and stochasticity. Data privacy is addressed by keeping the data processing mostly confined to on-computer tool and hook calls. The hook architecture allows for a variety of complicated imaging procedures, while simultaneously keeping the cost of processing low. It is relatively inexpensive for a model to build a single function or a script and, because we condition all code to be reusable both within and independently of MicroClaw, this is a one-time cost that produces a physical artifact with lasting value. We plan to further reduce costs by enabling MicroClaw to work with other LLMs, in particular local, open models. Local models could also be fine-tuned for microscopy tasks^33^, which we expect will improve performance. Due to model stochasticity, single MicroClaw sessions may decide to not use an available tool or skill, and some sessions use all information available in the persistent knowledge base while others ignore parts. However, the hooks and compiled scripts generated by MicroClaw are deterministic, so only one successful acquisition is needed to overcome this challenge. For training new users on a system, we still recommend an expert user be involved as imaging recommendations vary in quality.

An incomplete understanding of the microscope could in principle lead to poor hardware decisions, where a model could decide to move a stage into an objective or similar. We addressed this with a robust safety configuration that sits on top of tool calls and overrides MicroClaw’s decisions in the case an operation is unsafe. The user is guided to interact with and help properly set up the safety bounds in MicroClaw’s first launch session.

The capabilities demonstrated in this paper are just a few of many ways LLM-based agents can support microscopy. We plan to continue development of MicroClaw, adding a shared skills and hooks repository, new image analysis features–such as connections to cellpose^34^, µSAM^35^, and SMAP^36^, tools for writing custom GUIs, additional pre-loaded skills for light sheet and MINFLUX imaging, and additional support for OpenAI, Qwen^37^, Mistral^38^, and other LLMs. We plan to use our existing web-based GUI architecture to enable remote control of the microscopes, and enable MicroClaw instances on different machines to share memory^39^, allowing correlative experiments to be handled by MicroClaw across systems. We envision connections to other devices and software–such as enabling fully Python agentic microscopy control by connecting with pymmcore-plus^40^, smart, high-throughput SMLM workflows with fast imaging over a network in PYME^41^, and syncing spatial light modulators with acquisitions for fast MINFLUX tracking in Imspector^42^.

Today, MicroClaw is a microscopy agent that can augment and accelerate experiments. We believe MicroClaw will make previously challenging–and as of yet unthinkable– experiments into routine tasks. We believe this ushers in an exciting new era for biological science, where researchers can use microscopy agents to perform regular imaging and analysis tasks, and use the saved time and effort to pursue high-level experimental challenges and interesting imaging problems.

## Methods

### Agent-based design

MicroClaw was developed through iterative collaboration between the authors and coding agents Claude Sonnet 4.6, Opus 4.8, Opus 5, and Fable 5 (Anthropic, San Francisco, CA, USA), and Codex GPT-5.6-Sol and GPT-6-Astra (OpenAI, San Francisco, CA, USA), following an artifact-driven workflow^43^. Development was chunked by topic, where for each topic agents produced Markdown design plans, rationale, and implementation tasks. The authors reviewed and revised these documents, adding inline comments and follow-up prompts before implementation. Feedback was often phrased as questions to encourage critical evaluation rather than agreement^44^. Sessions were kept deliberately short, with the aim of limiting errors arising from exceeding the context window^45^, while persistent design documents supported continuity between sessions. A multi-agent workflow was adopted, where a coordinating agent assigned tasks to implementation agents, reviewed their code and test evidence, and integrated the results into the main code base. Validation included automated tests, tests on Micro-Manager running its demo configuration, and checks on physical microscopes. Findings informed subsequent code and design revisions. Recorded development prompts, design documents, implementation checklists, and validation records are available in the repository’s design directory.

### Software architecture

MicroClaw’s core functionality was kept as small as practical to support maintainability. The program runs an LLM agent that communicates with the user via a web graphical user interface and controls a running Micro-Manager instance through MicroClaw tools and Pycro-Manager’s ZMQ interface^12^, as shown in **Figure 1a** and **Supplementary Figure 4**. The web interface is designed to prevent unauthorized access: by default, it is available only on the microscope computer, and additional authentication and encryption safeguards are required for remote use. It also prevents multiple commands from controlling the microscope at the same time. Between the LLM and the microscope, safety and authorization gates enforce user-defined limits on stage motion, exposure, acquisition size and duration, illumination, channels, and device-property writes. Selected actions, such as enabling illumination or starting a large acquisition, require explicit user confirmation and are recorded for auditing. These safeguards depend on a correctly reviewed configuration, are not comprehensive collision-avoidance mechanisms, and do not replace user supervision.

MicroClaw ships with workflow skills distilled from expert knowledge and research literature. They cover areas such as authoring custom Python acquisition and analysis hooks, interpreting optical paths, operating particular focus and control systems, and acquiring SMLM and DNA-PAINT samples. A catalog of available skills is included in the agent’s permanent prompt, and the complete text of a relevant skill is loaded when the corresponding workflow is requested.

MicroClaw also maintains a machine-local knowledge base containing persistent information about the user’s microscope, samples, devices, calibrations, and preferred acquisition strategies. This information is supplied to the agent as reference data rather than executable instructions, and current hardware state must still be verified before it is used. User-directed additions require confirmation. The knowledge base is separate from the repository-maintained skills and allows MicroClaw to retain setup- and experiment-specific contexts across sessions.

Custom and package-specific acquisition analysis is integrated primarily through hooks and saved analysis adapters, while shared analysis primitives and several standard analyses remain in the main package. Hooks can observe images, record measurements, filter positions, influence adaptive acquisitions, generate guarded follow-up events, or connect external analysis packages to the acquisition. Hook source is shown to the user, checked for syntax and contract compatibility, reviewed with advisory security warnings, explicitly approved, and cryptographically pinned before later execution. These checks do not constitute a sandbox: hooks execute as Python code, and Micro-Manager plugin hooks may execute arbitrary Java outside MicroClaw’s normal safety checks. Optional analysis dependencies are installed only by users who need them, avoiding the need to impose every setup’s software stack on all installations. In cases where existing code or hooks have trouble interpreting an image, the `snap_and_analyze` tool can pass an image to the model for feedback.

### Microscopes

The custom SMLM setups for **Figures 2a, c**, and **d** were built following the design in ^19^, but using a 160x NA 1.43 oil-immersion objective (Leica)^20^. The effective pixel size was 127 nm.

**For Figure 2b**, origami fluorescence imaging was carried out on an inverted Nikon Eclipse Ti2 microscope (Nikon Instruments) with a Perfect Focus System, equipped with an Andor Dragonfly unit. The Dragonfly was used in the TIRF mode, applying an objective-type TIRF or HiLo configuration with an oil-immersion objective (Nikon Instruments, Apo SR TIRF 60×, NA 1.49, Oil). For excitation, a 561-nm laser was used. The beam was coupled into a multimode fiber going through the Andor Borealis unit reshaping the beam from a Gaussian profile to a homogenous flat top. As a dichroic mirror, a CR-DFLY-DMQD-01 was used. Fluorescence light was spectrally filtered with an emission filter (TR-DFLY-F600-050) and imaged with a scientific complementary metal oxide semiconductor (sCMOS) camera (Sona 4BV6X, Andor Technologies) without further magnification, resulting in an effective pixel size of 108 nm.

For **Figure 2b**, Exchange-PAINT, fluorescence imaging was carried out on an inverted Nikon Eclipse Ti1 microscope (Nikon Instruments) with a Perfect Focus System, equipped with a TIRF module, applying an objective-type TIRF or HiLo configuration with an oil-immersion objective (Nikon Instruments, Apo SR TIRF 60×, NA 1.49, Oil). For excitation, a 561-nm laser was used. As a dichroic mirror, a ZT 561 flat (Chroma) was used. Fluorescence light was spectrally filtered with an emission filter (600/50 ET Bandpass (Chroma)) and imaged with a scientific complementary metal oxide semiconductor (sCMOS) camera (Fusion BT, Hamamatsu Photonics) without further magnification, resulting in an effective pixel size of 108 nm. An Aladdin AL-4000 syringe pump (World Precision Instruments, Sarasota, FL, USA) was attached to this setup.

All systems were controlled via Micro-Manager software.

### Sample preparation

#### Clathrin

U-2 OS cells expressing endogenously SNAP tag-labelled clathrin light chains were grown at 37 °C, 5% CO_2_ in T25 cell culture flasks (Nunc) in phenol-free DMEM (Gibco) supplemented with 10% fetal bovine serum (Gibco), GlutaMAX (Gibco), non-essential amino acids (Gibco), and ZellShield ((Minerva Biolabs, 13-0050). To prepare samples for imaging, cells were grown on glass coverslips coated with Fibronectin (Bovine, 1 µg/cm^2^, Sigma-Aldrich) to 70% confluence. Cells were fixed in 3% PFA in cytoskeleton buffer (10 mM MES pH 6.1, 150 mM NaCl, 5 mM EGTA, 5 mM glucose, 5 mM MgCl_2_) for 12 min at room temperature, followed by quenching in 100 mM NH_4_Cl for 5 min. Cells were then permeabilized in 1% Triton-X for 10 min, followed by incubation in ImageIT FX signal enhancer (Invitrogen) for 30 min. Cells were then blocked in 2% BSA for 1 hr and stained using BG-AF647 (NEB) in 1% BSA, 0.1% Triton-X, and 1 mM DTT overnight. Cells were then washed in 0.1% Triton-X and PBS and stored at 4 °C until imaging.

#### DNA-origami and Exchange-PAINT

##### Buffers

Three buffers were used for sample preparation and imaging: Buffer A (10 mM Tris-HCl pH 7.5, 100 mM NaCl, 0.05% Tween 20, pH 7.5); Buffer B (10 mM MgCl2, 5 mM Tris-HCl pH 8.0, 1 mM EDTA, 0.05% Tween 20, pH 7.5), and Buffer C (1× PBS, 500 mM NaCl). The imaging buffers were supplemented with: 1× (+−)-6-hydroxy-2,5,7,8- tetra- methylchromane-2-carboxylic acid (Trolox), 1× 3,4-dihydroxybenzoic acid (PCA) and 1× protocatechuate 3,4-dioxygenase pseudomonas (PCD).

##### Trolox, PCA and PCD

100× Trolox: 100 mg Trolox, 430 μL 100% Methanol, 345 μL 1 M NaOH in 3.2 mL H_2_O. 40× PCA: 154 mg PCA, 10 mL water and NaOH were mixed, and pH was adjusted to 9.0. 100× PCD: 9.3 mg PCD, 13.3 mL of buffer (100 mM Tris-HCl pH 8.0, 50 mM KCl, 1 mM EDTA, 50% glycerol).

##### DNA origami self-assembly

All DNA origami structures were designed with the Picasso design tool^24^ (see **Supplementary Figure 2**). Self-assembly of DNA origami was accomplished in a one-pot reaction with 50 μL total volume, consisting of 10 nM scaffold strand (sequence see **Supplementary Table 1**), 100 nM folding staples (**Supplementary Tables 2-4**), 1 µM biotinylated staples (**Supplementary Table 7**), and 1 μM of docking site strands (for a list of DNA-PAINT handles see **Supplementary Table 5** and for imagers see **Supplementary Table 6**) in folding buffer (1× TE buffer (10 mM Tris and 1 mM EDTA) with 12.5 mM MgCl2). The reaction mix was then subjected to a thermal annealing ramp using a thermocycler. The reaction mix was first incubated at 80 °C for 5 min, then cooled from 60 to 4 °C in steps of 1 °C every 3.21 min, and then held at 4 °C.

##### DNA origami PEG purification

DNA origami structures featuring NanoLobster, a 10-nm and a 20-nm-grid were purified via two rounds of polyethylene glycol (PEG) precipitation by adding the same volume of PEG-buffer (15% PEG-8000, 500 mM NaCl in 1× TE buffer, pH 8.0), centrifuging at 14,000 g at 4 °C for 30 min, removing the supernatant, and resuspending in folding buffer.

##### DNA origami sample preparation

For chamber preparation, a piece of coverslip (no. 1.5, 18 × 18 mm^2^, ∼0.17 mm thick) and a glass slide (3 × 1 inch^2^ 1 mm thick) were sandwiched together by two strips of double-sided tape to form a flow chamber with an inner volume of ∼20 µL. First, 20 µL of biotin-labeled bovine albumin (1 mg/mL, dissolved in buffer A) was flown into the chamber and incubated for 2 min. Then the chamber was washed using 40 µL of buffer A. Second, 20 µL of streptavidin (0.5 mg/mL, dissolved in buffer A) was then flown through the chamber and incubated for 2 min. Next, the chamber was washed with 40 µL of buffer A and subsequently with 40 µL of buffer B. Then a mix of NanoLobster, 10-nm Grid and 20-nm Grid structures, each at a concentration of 100 pM, was incubated for 5 min. The chamber was then washed with 40 µl of buffer B again. Finally, the imaging buffer with buffer B and 1× Trolox, 1× PCA and 1× PCD with the Cy3B labeled imager (P3-Cy3B at 500 pM) strand was flown into the chamber. The chamber was sealed with Picodent before subsequent imaging.

##### Cell culture

U-2 OS cells were cultured in McCoy’s 5A Medium (Gibco) supplemented with 10% FBS (Gibco). The night before immunolabeling, cells were seeded on ibidi 8-well glass coverslips at ∼30,000 cells/well.

##### Plasmids

For labeling the outer membrane of mitochondria (**Figure 2b**), we expressed GFP-OMP25 from a plasmid as described before^46^. mCherry-Sec61β was acquired from Addgene (plasmid 49155).

##### Transient transfection

Transfections were performed using Lipofectamine 3000 (Invitrogen) according to the manufacturer’s instructions. Cells in eight-well glass-bottom chambers were transiently transfected with OMP25-GFP and Sec61b-mCherry and allowed to express for 16–24 hr before fixation.

##### Cell Fixation (Figure 2b)

Cells were fixed with 3% PFA and 0.1% GA for 15 min. After four washes (30 s, 60 s, 2× 5 min), cells were blocked and permeabilized with 3% BSA and 0.25% Triton X-100 at room temperature for 1 hr. Next, cells were incubated with primary antibodies against mCherry (anti-mCherry) and nanobodies against GFP (Massive Photonics) in 3% BSA and 0.1% Triton X-100 at 4 °C overnight. The next day, after four washes (30 s, 60 s, 2× 5 min), cells were incubated with secondary nanobodies (anti-Mouse) for 2 hr at room temperature. Finally, samples were washed three times with 1× PBS for 5 min each before adding the imaging solution.

#### Kinesin and Microtubules

U-2 OS derived stable cell lines were generated carrying a cassette, genomically integrated in the AAVS1 safe harbor locus, containing two constructs, Kif5B[1-560]-HaloTag7 and eGFP-ɑTubulin, under the control of the Tet-On® 3G inducible expression system and the CMV promoter, respectively. Cells were seeded on glass coverslips (Marienfeld, 117640) and grown for two days to favour the development of large lamellipodias. The expression of kinesins was induced by adding doxycycline at 10 µg/mL for 2 hr, followed by 30 min incubation with JFX646-HTL (Lavis Lab) added at 100 ng/mL. Then, cells were washed 3 times with fresh media, 10 min each wash, then taxol was added at 10 µM and incubated for 10 min before mounting for imaging with the same media containing taxol. At all times cells were cultured in the same media: DMEM (Gibco, 11880036) supplemented with 10% fetal bovine serum (Gibco, 10270106), GlutaMAX (Gibco, 35050038), non-essential amino acids (Gibco, 12084947), and ZellShield (Minerva Biolabs, 13-0050).

#### Mitochondria

U-2 OS cells (HTB-96™, ATCC) were cultured in McCoy’s 5 A medium with 10% FBS (Gibco #10270-106), 1 mM sodium pyruvate and phenol red (Gibco #16600082). Bacterial and fungal contamination was suppressed by addition of 1× ZellShield (Minerva Biolabs, 13-0050). The cells were grown in an incubator at 37 °C, 5% CO_2_ and 90% humidity. Before the cells reached confluency, they were split and seeded on plasma-cleaned 24 mm #1.5H round glass coverslips (Marienfeld) in a 6-well plate and left to adhere overnight. Apoptosis was induced using 10 µM ABT-737 (MedChemExpress #MCE-HY-50907-10mg) + 10 µM S63845 (MedChemExpress #MCE-HY-100741-5mg) in full medium. 20 µM QVD (MedChemExpress #MCE-HY-12305-10mg) was added to prevent the cells’ detachment from the coverslips.

Cells were fixed in 4% FA/PBS (Electron Microscopy Science #15710) at room temperature for 10 min, followed by quenching with 100 mM NH4Cl in PBS. After permeabilization in 0.5% Triton-X-100/PBS (Merck/Millipore #1086031000) for 5 min, cells were blocked for 1 hr in 5% BSA/PBS (Albumin (BSA) Fraction V, (pH 7.0), AppliChem #A13910500). For direct immunofluorescence the cells were incubated with an antibody against TOMM20 directly coupled to Alexa Fluor 647 (Abcam #ab209606, clone EPR15581-54) [for SMLM imaging] in 5% BSA/PBS for 1 hr at room temperature in a humidified chamber. The cells were then washed in PBS three times and incubated with another antibody against TOMM20 directly coupled to Alexa Fluor 488 (Abcam #ab205486, clone EPR15581-39) [for widefield imaging].

The coverslips were then mounted on custom stage holders, whose cavity was filled with ∼ 500 µL blinking buffer, final composition: [50 mM Tris-HCl pH 8.0, 10 mM NaCl, 10% [w/v] D-Glucose, ∼ 100 U/mL Glucose oxidase (GLOX), ∼ 2000 U/mL Catalase, 2.55 % [v/v] Glycerol, 35 mM MEA].

### Imaging and Analysis

#### Clathrin

Clathrin samples were imaged with a laser power of 5.8 kW/cm^2^. The SMAP software package^36^ served to process, reconstruct, drift-correct, and filter STORM data. A 10-nm localization precision filter and a 160-nm PSF filter were applied to all STORM data shown in **Figure 2a**.

#### DNA-origami and Exchange-PAINT

Imaging parameters for PAINT experiments are in **Supplementary Table 8**. Raw fluorescence microscopy images were subjected to spot-finding and subsequent super-resolution reconstruction, drift correction, filtering and alignment using the ‘Picasso’ software package^24^. x, y and z drift correction were performed with a redundant cross-correlation which is integrated in the same software package.

#### Kinesin and Microtubules

Kinesins were imaged with a laser power of 0.42 kW/cm^2^. Microtubules were imaged with a laser power of 0.23 kW/cm^2^. All kinesin analysis, including localization and tracking, was performed within MicroClaw using a custom tracking code similar to ^47^. Information is available in the history files and associated hook code.

#### Mitochondria

Mitochondria samples were imaged with a laser power of 11.2 kW/cm^2^. The widefield images were contrast adjusted in Fiji^48^. The 2D SMLM images were fitted with the simple fitting algorithm in SMAP^36^. The render settings were: display localizations as Gaussian of localization precision, set loc.prec <15, PSF <160, LLrel from -1 to 0 and contrast adjusted. Ilastik analysis was performed with the .ilp file provided in the data (see **Data availability**).

## Data availability

All data collected and the MicroClaw history files associated with them are stored in a Zenodo repository: https://doi.org/10.5281/zenodo.22768082.

## Code availability

Source code is available under a BSD-3-Clause license at https://github.com/Micro-Claw/microclaw. We welcome contributions from the community.

## Acknowledgements

We thank Pallavi Deolal and Phylicia Kidd for helpful discussions. This work was supported by the Austrian Science Fund (10.55776/ESP3754525 to ZM, PAT1965125 to JR) and the European Research Council (MSCA 101201217 to SVS, PoC 101199020 to JR). FS acknowledges funding from EPFL for the Laboratory of Molecular Spatial Omics.

## Author Information

ZM, FS and JR conceived of the project. ZM, AA, NC, SVS, FH, LTK, FS and JR designed the software. Coding agents implemented the software, with the help of ZM, FH and NS. AA, NC, SVS, LTK, AM, and FS prepared and imaged samples. ZM also imaged samples. ZM, FS and JR analyzed the results and wrote the paper with help from all of the authors.

## Competing Interests

The authors declare no competing interests.

## Supplementary Information

**Supplementary Figure 1:**
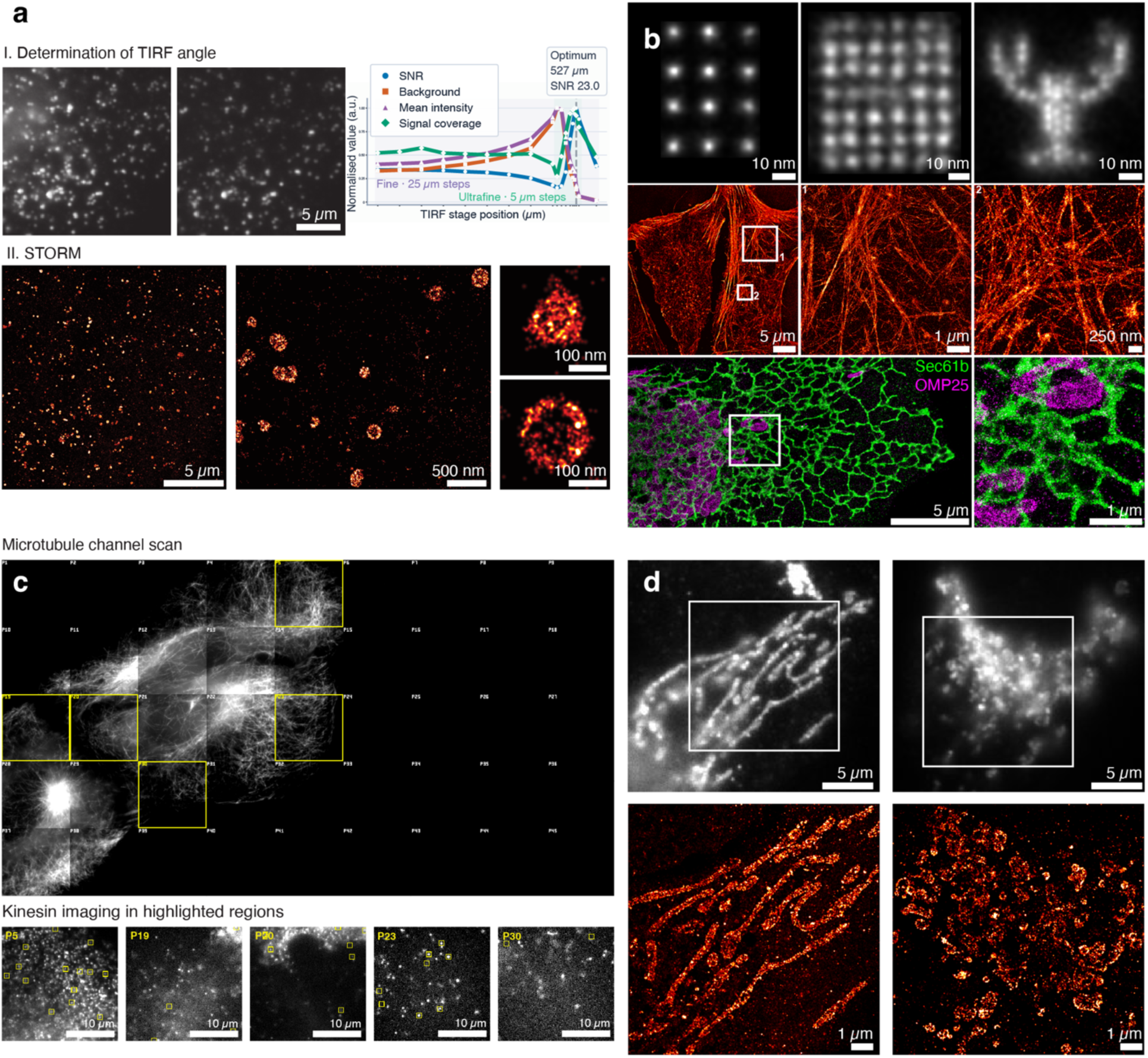
Images taken using the MicroClaw workflows in Figure 2. **a,** TIRF angle optimization (**i**) of a clathrin sample, followed by SMLM imaging (**ii**). Performed on custom SMLM system^19^. The agent optimizes TIRF with four metrics (**i**, right) to move from epi illumination (**i**, left) to TIRF (**i**, middle). In the resulting TIRF SMLM images, spherical clathrin pits are partially sampled as rings, while flat lattices stay in focus, as expected. **b,** PAINT imaging of DNA-origami (see **Supplementary Figure 2**) on a Nikon-Eclipse Ti (grids) and Andor Dragonfly (lobster, see **Supplementary Figure 3**) systems. We successfully resolve 20 nm and 10 nm grids and 5 nm spacing on the lobster, with 0.9 angstrom precision. Exchange-PAINT of cells taken on a Nikon-Eclipse Ti. Here the agent informed us when to push in new imager strands with the syringe. MicroClaw handled the rest. PAINT imaging of actin (red). DNA-PAINT imaging of ER (green) and mitochondria (magenta). **c,** Detection of walking kinesin on a coverslip. Microclaw scanning large sample areas of a mixture of live U-2 OS cells expressing either Nup96-GFP or Kinesin-HaloTag7-JF646, capable of distinguishing and focusing on the microtubules, then autonomously switching to a second channel to identify walking kinesins, their density and motility. **d,** Healthy vs. apoptotic mitochondria detection. Here we connect MicroClaw to the external segmentation program, Ilastik^14^, and use a pre-trained model to detect mitochondrial structures in regions of interest in a tiled acquisition and determine if they are apoptotic or healthy. One of each is shown imaged in super-resolution.

**Supplementary Figure 2:**
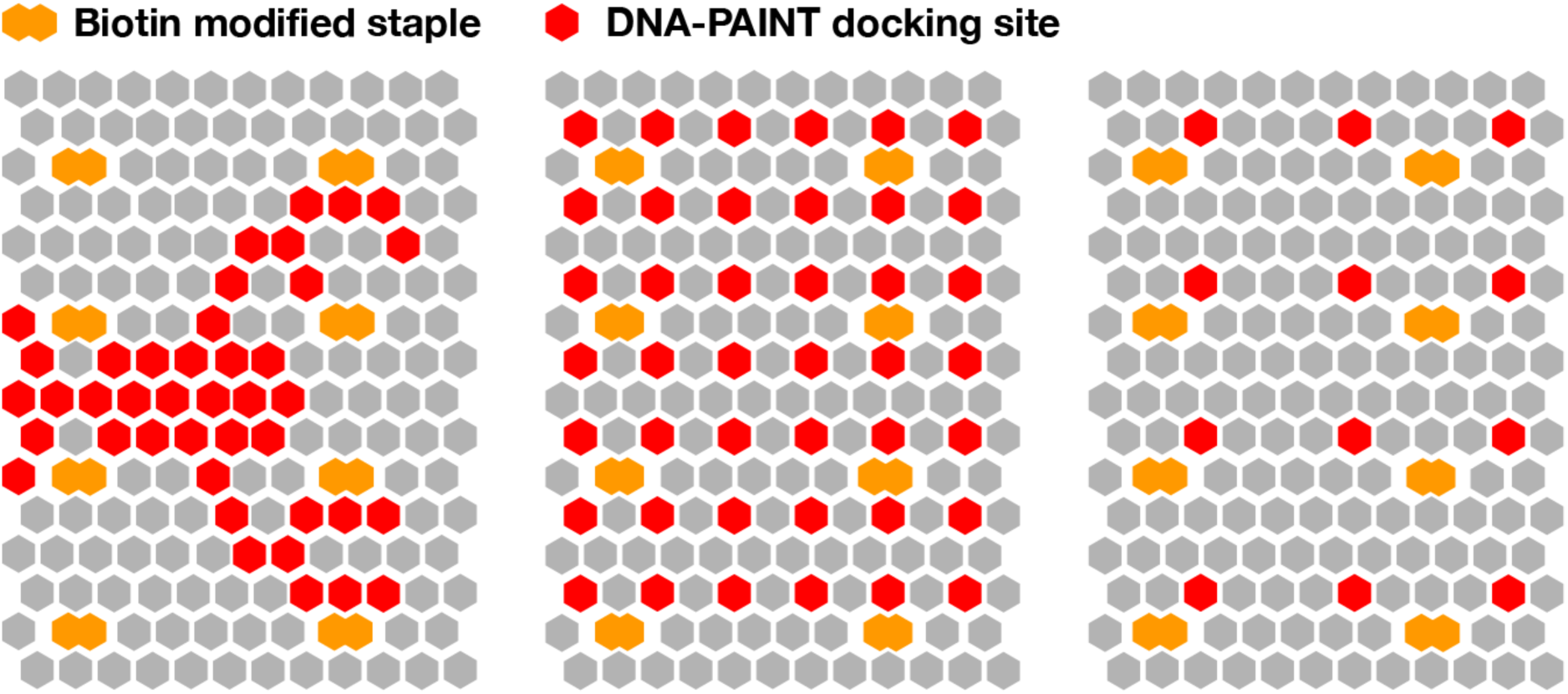
Used DNA origami designs. Schematic representation of all DNA origami designs used in this study. The hexagons represent 3’-staple positions. Red hexagons are representing staples extended with docking sites for transient binding of Imager probes. The orange hexagons depict staples extended with a biotin modification for immobilization on the cover slip surface.

**Supplementary Figure 3:**
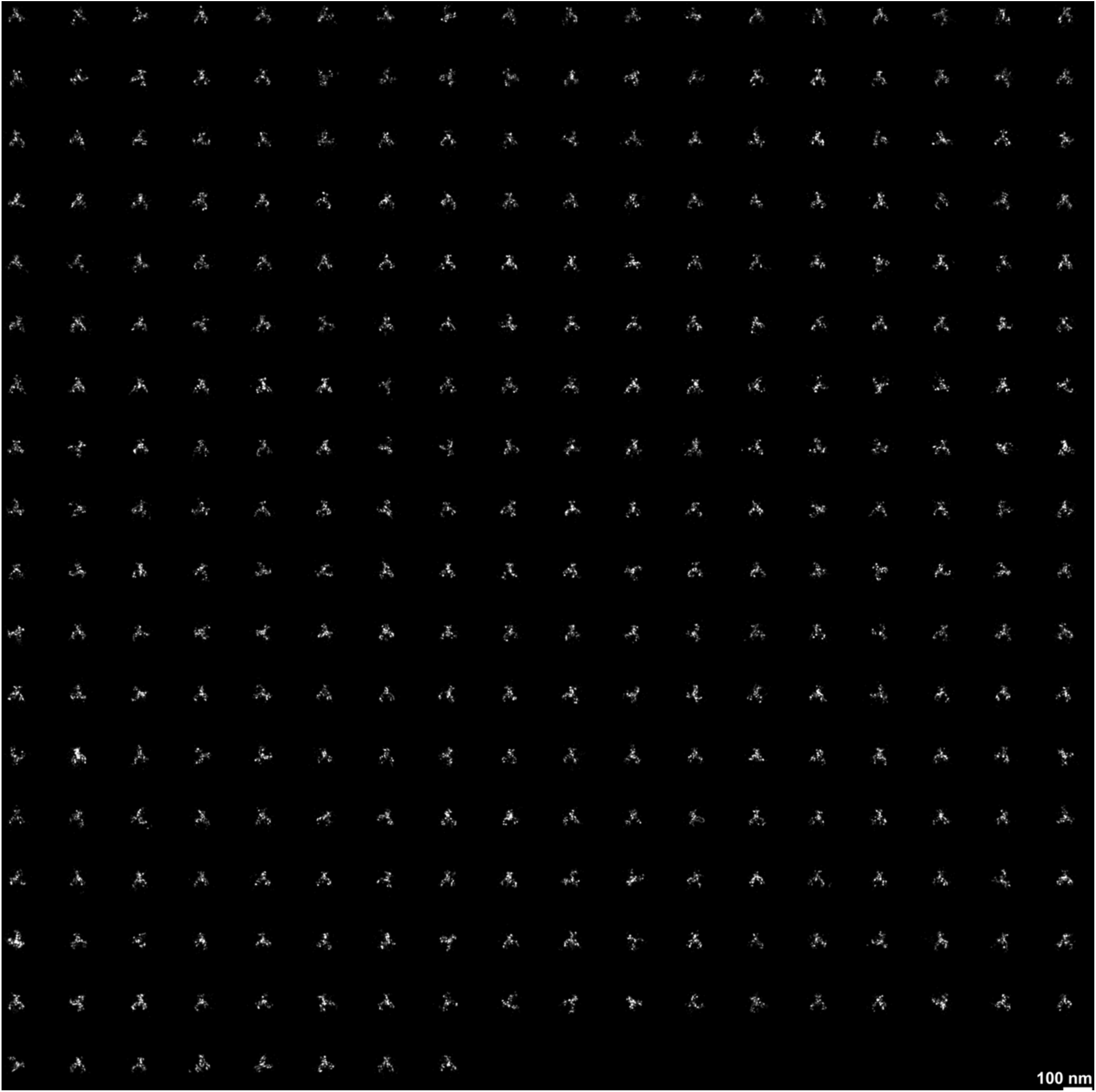
Single DNA origami structures displaying the NanoLobster. Images of the 314 single structures used to produce the averaged image depicted in **Supplementary Figure 1b.**

**Supplementary Figure 4:**
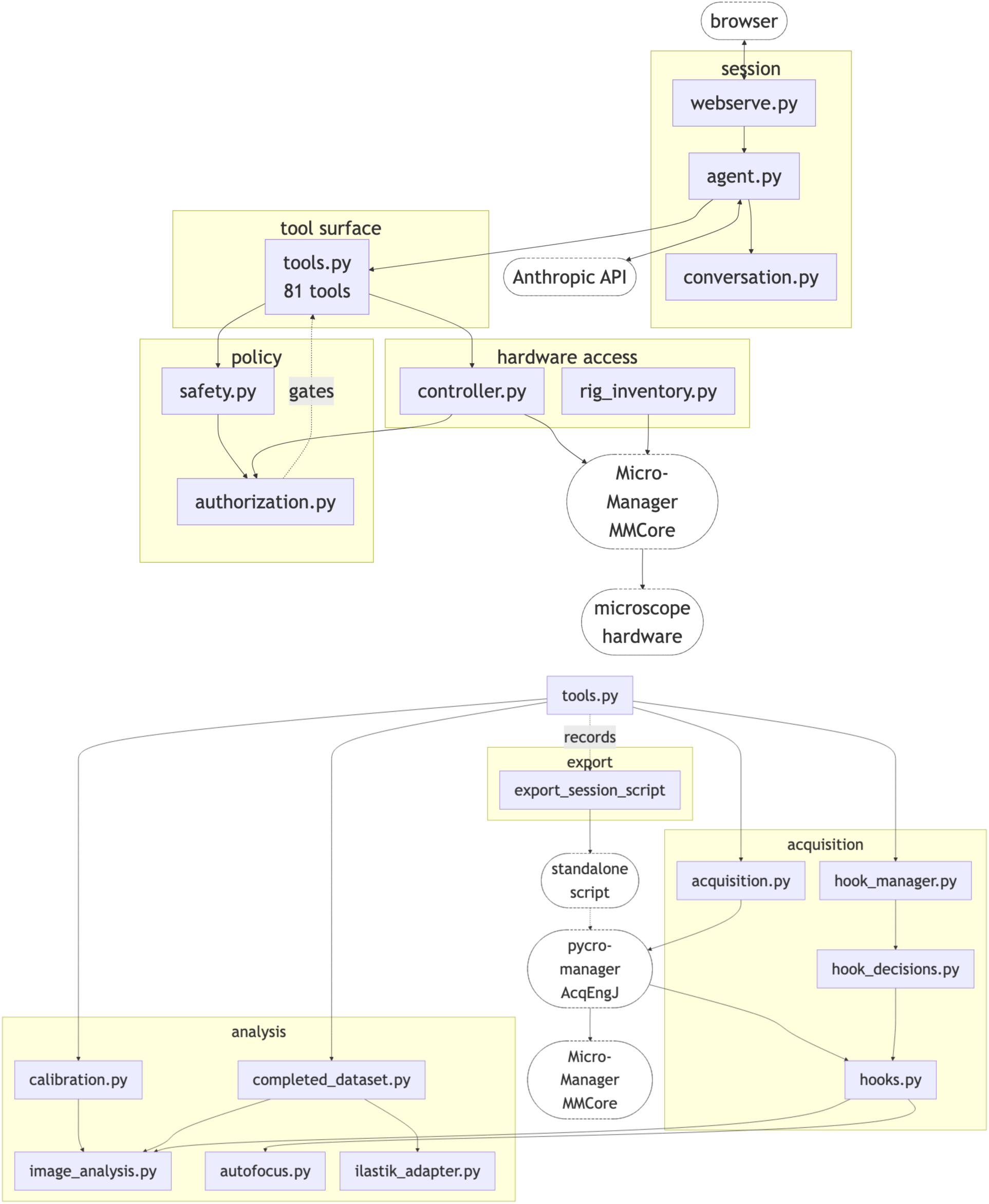
Mermaid diagram of MicroClaw. This shows the full schematic of interactions of various parts of the agent and its external elements.

## Supplementary Note 1

This includes a summarized transcript of a conversation with MicroClaw to perform a simple task: autofocusing and then recording, done via two separate paths: an internal autofocusing tool and Micro-Manager’s OughtaFocus plugin. Repeated explanations and raw tool payloads are condensed; significant tool results are incorporated. Editorial notes distinguish inconsistencies or unverified claims in the original from recorded results. The full transcript and the resulting script can be found in the data (see **Data availability**).

### Conversation

**User**: I have beads on a coverslip. Using the current settings, take a 3×3 tile scan with autofocus at each position. Tell me whether the beads are approximately in the same focal plane and how many beads are at each position.

**MicroClaw**: I checked the microscope. It is using Res1, 0.127 µm/pixel, a 233×206-pixel ROI, and 50 ms exposure. The 638 laser is already enabled at about 2%; the other lasers are off. Hardware focus lock is disengaged, with QPD X railed at 65535. I will use software autofocus and leave the existing illumination settings in place.

I propose one saved frame at each of nine positions, a 26 µm tile step, and an initial autofocus range of 20 µm with 0.5 µm steps around the current Z of approximately 54 µm. I will read the focus log and count beads using bead_counter_v2. These will be unvalidated blob/component counts, although the saved method was previously compared with visual counts. Please confirm the current excitation settings and a dated folder under F:\DataSSD.

**User**: Yes. The folder name sounds good; add my initials, ZM.

**MicroClaw**: I will save to F:\DataSSD\20260910_ZM_beads_3x3_autofocus.

The first acquisition was blocked: the autofocus safety check predicted Z=104.001 µm, above the 100 µm limit. Current Z read back as 54.003 µm. Reducing the requested range to 12 µm was also blocked, with a predicted maximum of 101.999 µm. The position list was empty, and a test position at Z=54 µm passed validation. I offered several possible explanations involving grid placement and autofocus search arithmetic, but did not establish the cause. I reduced the range to 6 µm, retaining 0.5 µm steps.

That run completed: nine positions, one saved image each, and autofocus marked converged at every position. The tool recorded 121 autofocus exposures in addition to the nine saved frames. The dataset is beads_3x3_1, with autofocus_log.json in the parent folder.

The focus results and bead counts are:

| Tile | Focus Z ($\mu\text{m}$)* | Bead count |
| --- | --- | --- |
| r0_c0 | 56.002 | 2 |
| r0_c1 | 56.506 | 4 |
| r0_c2 | 56.514 | 2 |
| r1_c0 | 56.016 | 8 |
| r1_c1 | 56.514 | 7 |
| r1_c2 | 56.016 | 2 |
| r2_c0 | 56.016 | 6 |
| r2_c1 | 56.014 | 4 |
| r2_c2 | 56.018 | 4 |
| Total |  | 39 |

Tile Focus Z (µm)* Bead count

I concluded that the beads were approximately in the same focal plane: the Z spread was about 0.51 µm, comparable to the 0.5 µm autofocus step. Counting used background-subtracted LoG blob detection with threshold 150 ADU, minimum peak above background 300 ADU, and sigma range 1–4. Closely spaced beads can merge into one detection; these are screening counts, not a certified particle total. No detection-overlay image was produced. I can open the dataset for visual checking.

**User**: Yes, show me the dataset so I can verify the counts.

**MicroClaw**: I opened beads_3x3_NDTiffStack.tif in the Micro-Manager/ImageJ viewer: 233×206 pixels, nine planes. Planes 1–9 correspond to r0_c0, r0_c1, r0_c2, r1_c0, r1_c1, r1_c2, r2_c0, r2_c1, and r2_c2. Adjust contrast per plane to see dim beads. If a count looks wrong, I can adjust the detection threshold or prominence floor.

**User**: Counts look good! Repeat the procedure with Micro-Manager’s OughtaFocus plugin. I moved the beads about 2 µm out of focus again just for fun.

**MicroClaw**: OughtaFocus is installed. I can use the autofocus_mm_plugin hook with plugin_name=’OughtaFocus’. The plugin controls its own Z search; Microclaw checks the final position but does not guard the plugin’s motion during the search. Its search settings come from its own GUI configuration. Shall I read those settings first?

**User**: Yes.

**MicroClaw**: My attempt to read OughtaFocus through device properties failed because it is a plugin rather than a loaded hardware device. The available interfaces did not let me read its search range or metric. Please check its GUI settings; a range around 6–10 µm was my suggestion for this starting point. Focus lock remains off.

**User**: Where can I find OughtaFocus in the MM GUI?

**MicroClaw**: Use the gear/wrench beside the autofocus controls in the main Micro-Manager window, then select OughtaFocus in the method dropdown. I also suggested a Plugins > Autofocus route, depending on the build. Check SearchRange_um, Tolerance_um, the Maximize metric, and exposure; the plugin’s focus exposure may differ from the saved-image exposure.

**User**: Route 1 worked. Search range is 10 µm. Should be good. Let’s try it!

**MicroClaw**: I repeated the 3×3 scan with 26 µm steps and 50 ms saved-image exposures, using OughtaFocus at each position. All nine images were acquired and saved to beads_3x3_oughtafocus_1 in the same parent folder.

The acquisition initially reported all nine autofocus outcomes as ‘unknown’. I first said a focus table might not be available, but reading oughtafocus_log.json showed a best-Z value for every tile. There was still no explicit convergence verdict.

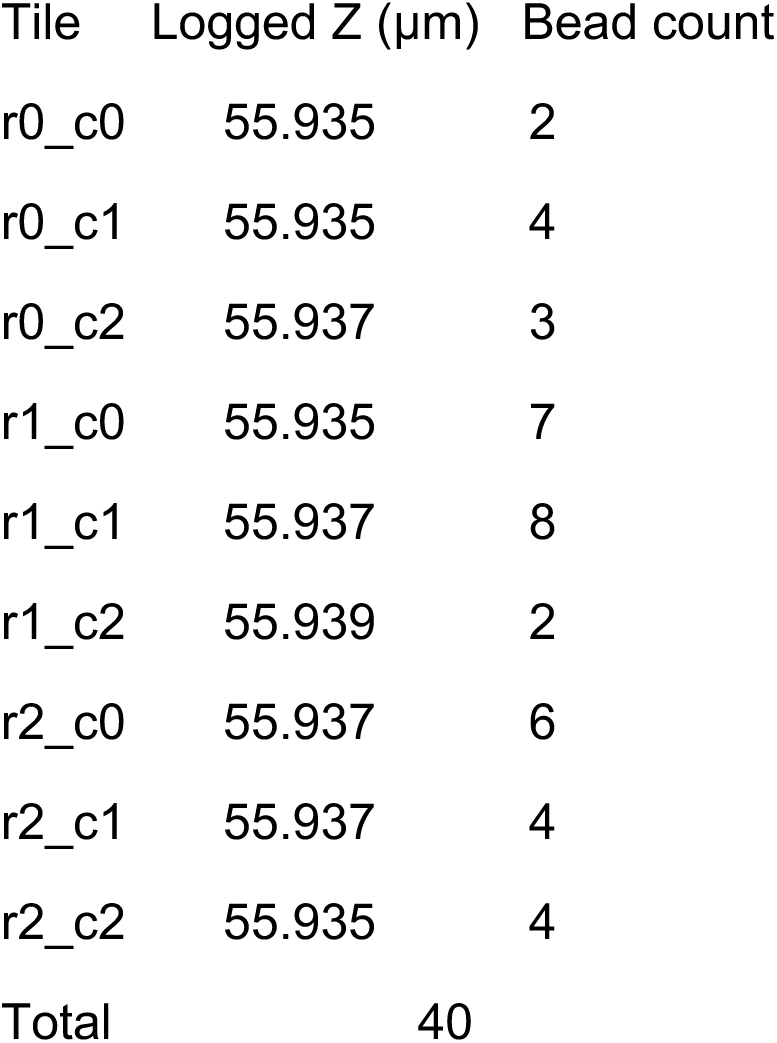

I again concluded that the beads were approximately in one focal plane. The recorded Z values span 55.935–55.939 µm. I reported OughtaFocus settings of SearchRange 10 µm, tolerance 1 µm, metric ‘Edges’, and 100 ms focus-search exposure, while saved frames used 50 ms. Bead counting used the same parameters as before.

Six tiles have identical counts between runs. Three differ by one: r0_c2 changes 2→3, r1_c0 changes 8→7, and r1_c1 changes 7→8. I suggested borderline detections and slight focus/frame differences as possible explanations, without testing the cause.

**User**: Yes, open that dataset too.

**MicroClaw**: I opened beads_3x3_oughtafocus_NDTiffStack.tif, again a 233×206-pixel stack with nine planes in the same tile order. Planes 3, 4, and 5 contain the changed counts. Contrast adjustment can help with dim beads. The 638 laser remains on, as it was at the start; would you like it left on or shuttered?

**User**: Again looks good. Could you export this session to a script so I can rerun these?

**MicroClaw**: I exported beads_3x3_session.py. The export tool returned complete=true, no not-emitted calls, and two emitted calls corresponding to the successful tile acquisitions. It explicitly skipped the two failed acquisitions. A file search confirmed the script exists and is about 207 KB.

I interpreted this as an export containing the scans but omitting bead counting, focus-log readouts, and viewer operations, and described it as an export capability gap. I offered a separate, handwritten companion driver for both scans and counting, warning that it would not be run or validated. Do you want counting included, and should the grid be fixed to the original coordinates or centered on the current stage position?

## Supplementary Note 2

This includes a summarized transcript of a conversation with MicroClaw to identify the source of a double image on the microscope’s camera. Repeated updates and raw tool output have been condensed into the dialogue. Image observations and optical explanations below reflect MicroClaw’s statements in the original conversation, including its initial diagnosis and subsequent correction; they are not an independent verification. The full transcript and the resulting script can be found in the data (see **Data availability**).

### Conversation

**User:** Hi! I see two copies of the bead image, and the beads are oddly shaped. Why? I expect one image with a symmetric point spread function (PSF).

**MicroClaw:** I’ll check the rig’s optical path. Servos.Position1 is 28000; your knowledge base identifies this as the 3D/astigmatic optic, with 4000 meaning out. Thorlabs ELL9 is at Position 1, while ELL9-1 is at Position 0; your recorded single-channel (1C) configuration has both sliders at Position 0. My initial suspicion is that the ELL9 slider causes the doubling and the astigmatic optic causes the asymmetric shape. Would you like me to restore the plain 1C configuration, or is the split/astigmatism intentional? The 638 nm laser is also enabled at 100%; I understand it to be camera-triggered, firing during exposures.

**User:** Yes, I want a plain, symmetric 1C PSF. The laser power seems awfully high. Let’s try 2%.

**MicroClaw:** I’ve set Servos.Position1 to 4000 (astigmatic optic out), moved Thorlabs ELL9 to Position 0, and set the Luxx638 laser power to 2%. The optics writes succeeded; the subsequent system readback shows both ELL9 sliders at Position 0 and the laser at about 2.0% (2.0024%). I concluded that the slider caused the doubling and the astigmatic element caused the distorted shape, so I expected a single symmetric PSF. The 638 laser remains enabled. Would you like live view started, or should I leave things as they are?

**User:** I still see two copies of the beads.

**MicroClaw:** Then the ELL9 slider I moved did not account for the remaining split. I checked other path elements and took a snapshot. Each bead appears round now, but has a companion spot at a fixed horizontal offset to its left: the whole field is duplicated. The snapshot has an SNR of 8.83 and no saturation at 50 ms exposure.

Servos.Position0 is 10000, the recorded empty 1C position for the transmitted/reflected slider. Servos.Position3 is 32000, identified in your knowledge base as the 700/100 far-red emission filter. ELL9-1 is at Position 0, recorded as the BFP/Bertrand lens out. I interpreted these motorized settings as correct for plain 1C imaging and suspected a manual prism, wedge, or two-channel image splitter that Micro-Manager cannot see. Removing the astigmatic optic appears to have fixed the shape; the duplication is a separate problem. Is there a manual splitter on the scope? I can consult the optical-path guidance. If you are going to touch the optics, tell me so I can stop the 638 laser and pause exposures.

**User:** Maybe. Where can I look for a manual splitter?

**MicroClaw:** I consulted the optical-path guidance, which emphasizes that Micro-Manager reports motorized components but cannot report manual prisms, sliders, and other physical components. Look first between the microscope port and the Andor camera for an image-splitter box, such as an OptoSplit, DV2, or W-VIEW, with a 1C/2C selector, bypass slider, flip mirror, or removable splitting cube. Other candidates are a manual port-selector prism, a filter cube with a beamsplitter or wedge, a dual-band emission filter, or a pellicle/wedge in a manual mount on the emission bench. Given the uniform horizontal offset, my strongest suspicion is a camera-port image splitter left in split mode. Tell me before reaching in so I can stop the laser and avoid exposures.

**User:** This is a custom system, but I can look in front of the camera. Do you think it’s before or after anything else?

**MicroClaw:** I interpreted the uniform horizontal displacement as evidence for a splitter acting on the formed image after the tube lens, immediately in front of the camera. I said a splitter earlier in the path would typically produce differences in focus, magnification, or separation across the field, and therefore suggested looking at the last optical components before the sensor. Look for a slit or entrance aperture, a beamsplitter plate/cube and fold mirror, a single-pass/split selector, a removable splitting cube, or a wedge/pellicle that can be bypassed. The 638 laser remains enabled and, as I described it, camera-triggered. I am not exposing right now; tell me before handling the optics so I can prevent an exposure.

**User:** There was a dichroic mirror before the two filter wheels in front of the camera. I took it out. Was that it?

**MicroClaw:** I took another snapshot and reported that the doubling was gone: each bead was now a single symmetric round spot, and the populated field was roughly half as wide. I attributed the two shifted copies to the manual dichroic before the camera’s two filter wheels. My final conclusion was that removing the astigmatic optic fixed the unusual PSF shape, while removing the manual dichroic fixed the doubling. The new snapshot had an SNR of 7.6 and no saturation, and I described the signal quality as comparable.

The 638 laser remains enabled at 2%, described as camera-triggered. Would you like me to save a knowledge-base note about this manual dichroic causing horizontal image doubling, and open live view?

## Notes

https://doi.org/10.5281/zenodo.22768082

https://github.com/Micro-Claw/microclaw

### Competing Interest Statement

The authors have declared no competing interest.

